# Lamellar cells in Pacinian and Meissner corpuscles are touch sensors

**DOI:** 10.1101/2020.08.24.265231

**Authors:** Yury A. Nikolaev, Viktor V. Feketa, Evan O. Anderson, Elena O. Gracheva, Sviatoslav N. Bagriantsev

## Abstract

The skin covering the human palm and other specialized tactile organs contains a high density of mechanosensory corpuscles tuned to detect transient pressure and vibration. These corpuscles comprise a sensory afferent neuron surrounded by lamellar cells^1-3^. The neuronal afferent is thought to be the mechanical sensor within the corpuscle, whereas the function of lamellar cells is unknown^2,4,5^. Here we show that lamellar cells within Meissner and Pacinian corpuscles detect tactile stimuli. We develop a preparation of bill skin from tactile-specialist ducks that permits electrophysiological recordings from lamellar cells and demonstrate that they contain mechanically-gated ion channels. We also show that lamellar cells from Meissner corpuscles generate mechanically-evoked action potentials using R-type voltage-gated calcium channels. These findings provide the first evidence for R-type channel-dependent action potentials in non-neuronal cells and demonstrate that lamellar cells are active detectors of touch. We propose that Meissner and Pacinian corpuscles use both neuronal and non-neuronal mechanoreception to detect mechanical signals.

---

The sense of touch is essential for a range of physiological processes, including detection of pain and pleasure, object recognition, foraging, and environment navigation. It facilitates the establishment of maternal bonds and underlies the development of social behaviors ^6^. The human palm contains a dense population of mechanosensory corpuscles that are tuned to detect transient pressure and vibration. Corpuscles are thus essential for precise manipulation of tools and objects, and performing fine tactile tasks ^1-3^. Animals that are mechanosensory specialists possess organs that are functionally analogous to the human palm, including the star organ of the star-nosed mole and the bill of tactile-foraging waterfowl. These organs contain hundreds of corpuscles per square millimeter of skin, allowing mechanosensory specialists to rely on touch during their search for food ^7-10^.

The two most common corpuscles in vertebrates are layered (Pacinian) and non-layered (Meissner) corpuscles. Layered corpuscles detect high-frequency vibration, whereas non-layered are tuned to lower frequencies ^3,7,11^. Both types are innervated by myelinated mechanoreceptors that arise from somatosensory ganglia. Neuronal mechanoreceptors are thought to be the only touch sensors within corpuscles and produce rapidly-adapting firing patterns when their mechanically-gated ion channels are activated by touch ^2,4,5^. In layered corpuscles, the mechanoreceptor is surrounded by onion-like sheaths formed by lamellar cells, whereas it is sandwiched between two or more lamellar cells in non-layered corpuscles. The functional role of lamellar cells is obscure, but they are thought to provide structural support for the neuronal afferent, facilitate small-amplitude vibrations ^12^ and serve as a passive mechanical filter for static stimuli ^13^. Interestingly, there are reports that some lamellar cells are immunoreactive for synaptic proteins, suggesting an active, rather than passive role in touch sensing ^14-16^. However, despite their widespread presence in vertebrates, the biophysical properties and physiological roles of lamellar cells remain unknown ^15^.

To test whether lamellar cells play active role in the detection of touch, we developed a glabrous skin preparation from the bill of Pekin duck, a tactile specialist bird ^7,17^. Duck bill skin contains a dense population of Pacinian and Meissner corpuscles, referred to as Herbst and Grandry corpuscles, respectively ^18,19^. Like their mammalian counterparts, duck corpuscles are innervated by rapidly-adapting mechanoreceptors and are tuned to detect transient pressure and vibration ^19-22^. Optical and electron microscopic analyses of an *ex vivo* preparation of duck bill skin (Fig. 1A and *Materials and Methods*) revealed a mixed population of Pacinian and Meissner corpuscles, which could be distinguished by their unique morphology and size (Fig. 1B and C). Duck Pacinian corpuscles had an oval structure, ∼35-120 µm in size (n=140 corpuscles), and comprised a mechanoreceptive neuronal afferent surrounded by an inner core and outer capsule formed by lamellar cells (Fig. 1D-F). Meissner corpuscles were spherical and smaller in size (∼15-35 µm in diameter, n=50 corpuscles) than Pacinian corpuscles. They consisted of a neuronal mechanoreceptor surrounded by two or more lamellar cells (Fig. 1G-I) ^14,18^. The presence of both types of corpuscle in duck bill skin suggests it is a good model system for the human palm, in contrast to mouse glabrous skin, which normally lacks layered corpuscles ^23^.

**Fig. 1.**
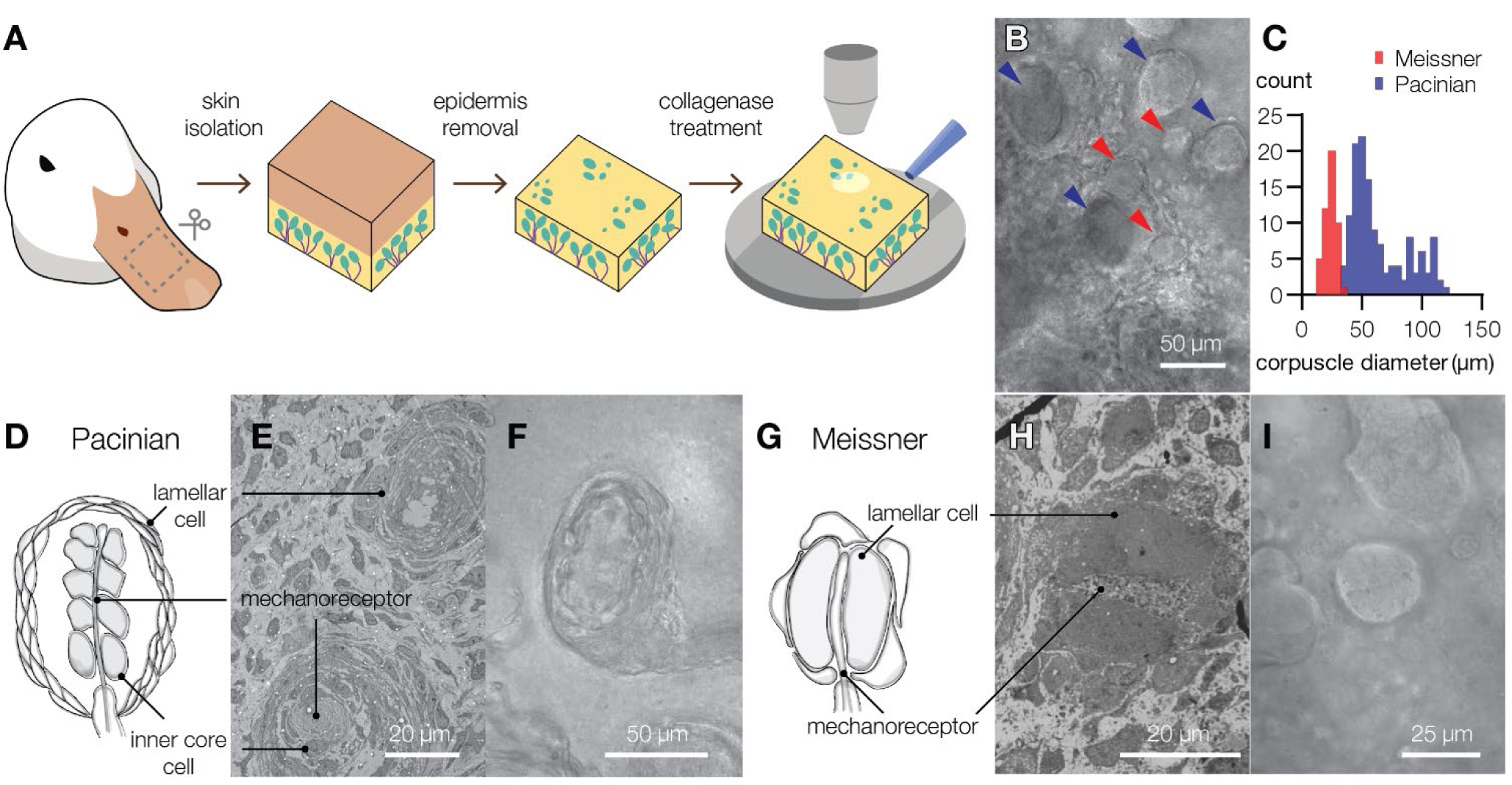
The bill skin of a tactile specialist duck possesses Pacinian and Meissner corpuscles. **(A)** Schematic illustration of the preparation of duck bill skin for electrophysiological and optical analysis of mechanosensory corpuscles. **(B)** A bright field microscopic image of a mixed population of Pacinian corpuscles (blue arrowheads) and Meissner corpuscles (pink arrowhead) in a patch of duck skin from the dorsal surface of the upper bill. **(C)** Size distribution of visible Meissner and Pacinian corpuscles in duck bill skin (50 Meissner and 140 Pacinian corpuscles total). **(D-I)** Illustrations (*D, G*), electron microscopy images (*E, H*) and close-up bright field microscopy images (*F, I*) of mechanosensory corpuscles. Pacinian corpuscles are composed of outer core lamellar cells surrounding an inner bulb of inner core cells and a neuronal mechanoreceptor. In Meissner corpuscles, the mechanoreceptor is sandwiched between two or more lamellar cells.

Having identified lamellar cells in mechanosensory corpuscles from duck bill skin, we sought to characterize them *in situ* by injecting the fluorescent dye Lucifer yellow using a patch pipette (Fig. 2A and B). The dye remained confined within the volume of each cell for 15 minutes post-injection, suggesting that a diffusion barrier existed between lamellar cells in both corpuscular types. The long, flat outer lamellar cells in Pacinian corpuscles had an average length of 13.5 ± 0.3 µm (mean ± s.e.m., n=5 cells, Fig. 2A). The hemi-spherical lamellar cells in Meissner corpuscles had an average diameter of 15.7 ± 1.4 µm (n=4 cells, Fig. 2B). Electrophysiological recordings revealed that Pacinian and Meissner lamellar cells had a whole-cell membrane capacitance of 9.6 ± 1.4 pF and 24.6 ± 4.6 pF, respectively (Fig. 2C). In addition, Pacinian lamellar cells had a resting membrane potential of −51.9 ± 2.0 mV and a high apparent input resistance of 5.8 ± 1.8 GΩ, whereas Meissner lamellar cells had a significantly more negative resting potential of −73.5 ± 2.4 mV and lower input resistance of 1.5 ± 0.4 GΩ (Fig. 2C).

**Fig. 2.**
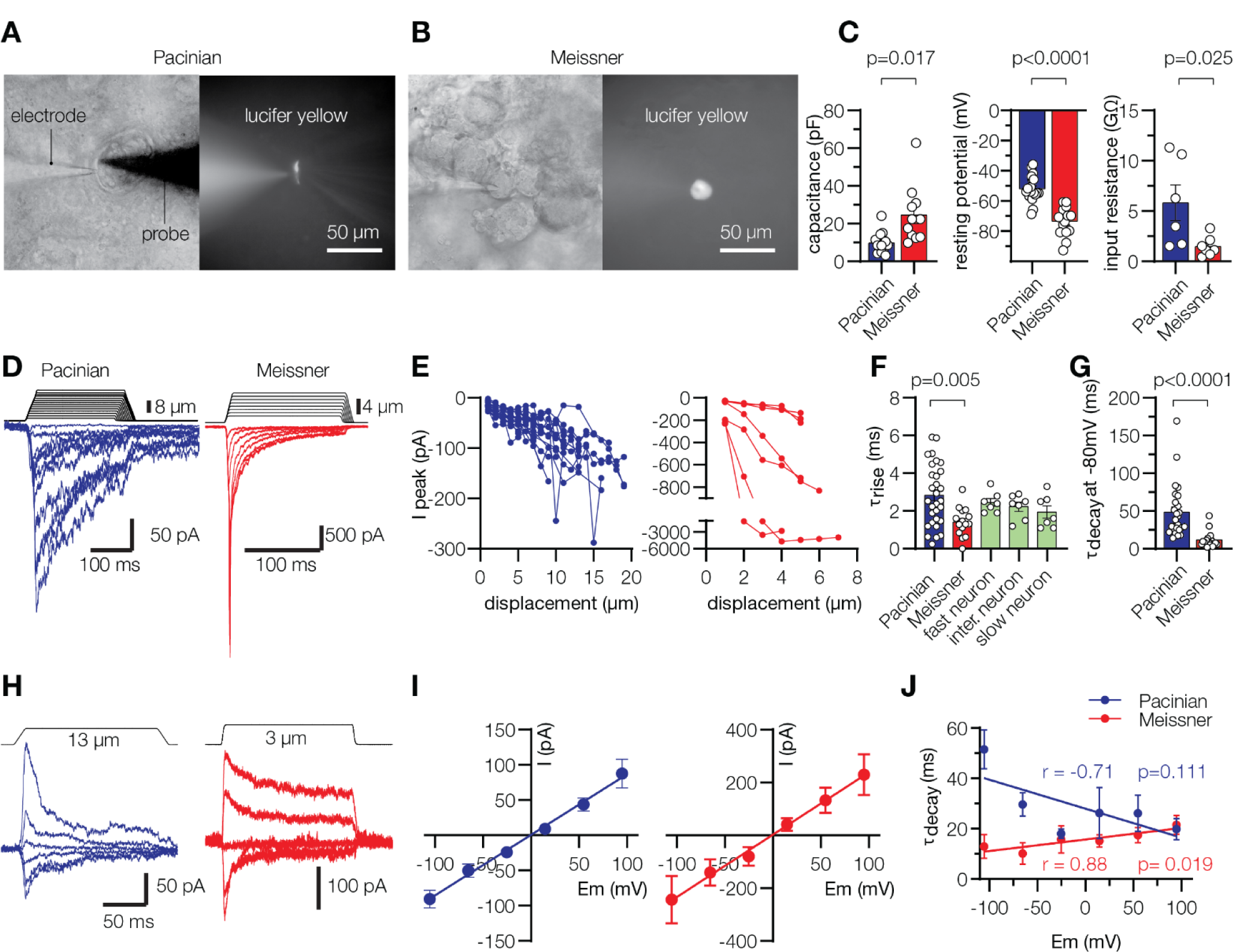
Lamellar cells of Pacinian and Meissner corpuscles are mechanosensitive. **(A, B)** Representative images of lamellar cells from Pacinian and Meissner corpuscles filled with Lucifer yellow via the recording electrode. A glass probe is positioned nearby to deliver mechanical stimulation. **(C)** Electrophysiological characteristics of lamellar cells. Significance calculated using unpaired two-tailed *t*-test. **(D)** Representative MA currents elicited from lamellar cells by mechanical indentation using a glass probe. **(E)** Quantification of peak MA current amplitude in Pacinian (*left*, n=19 cells) and Meissner (*right*, n=6 cells) lamellar cells in response to indentation with a glass probe. Lines connect measurements from individual cells. **(F)** Quantification of MA current rise time (τ_rise_) recorded in lamellar cells, and in trigeminal mechanoreceptors with fast, intermediate and slow MA current. The effect of treatment is significant, *F*_4,61_=3.49, p=0.013, one-way ANOVA with Tukey’s post-hoc test. **(G)** Quantification of lamellar cell MA current inactivation rate (τ_inact_). Significance calculated using two-tailed Mann-Whitney *U*-test (*U*=29).**(H)** Representative MA currents elicited from lamellar cells in response to indentation at different voltages. **(I)** Voltage-dependence of peak MA current from 8 Pacinian and 5 Meissner lamellar cells, fitted to the linear equation. **(J)** Quantification of MA current τ_inact_ from 7 Pacinian and 7 Meissner lamellar cells, fitted to the linear equation. r, Pearson’s correlation coefficient; p, probability of the line slope = 0. Data are presented as mean ± s.e.m. from at least three independent skin preparations. Open circles denote individual cells.

We next asked whether lamellar cells are mechanosensitive *in situ*. Stimulation of either Pacinian or Meissner lamellar cells with a glass probe produced robust mechanically activated (MA) currents, which increased in amplitude as probe displacement increased (Fig. 2D and E). Although MA currents from Pacinian lamellar cells had a significantly slower rise time than Meissner cell currents (τ_rise_ = 2.8 ± 0.3 ms and 1.4 ± 0.2 ms for Pacinian and Meissner cells, respectively, p=0.005), both values were within the range of MA currents recorded from mechanosensitive neurons (Fig. 2F) ^24,25^. Following activation, Pacinian lamellar MA currents decayed (τ_decay_ = 48.7 ± 7.0 ms), reaching 20%-68% of their peak amplitude by the end of the 150 ms stimulus (Fig. 2D and Extended Data Fig. 1A and B). In some cells, up to 30% fraction of peak MA current persisted after retraction of the probe, and in each case returned to baseline within 10 s (Extended Data Fig. 1C). In contrast, Meissner lamellar MA currents decayed significantly faster (τ_decay_ = 11.8 ± 2.3 ms, p<0.0001), and lacked a persistent, non-inactivating component (Fig. 2D and G). Both types of MA current had a linear voltage dependence and a near-zero reversal potential (Fig. 2H and I), characteristic of a non-selective cation conductance. However, they differed in their voltage dependence of inactivation; depolarization slightly decreasing τ_decay_ in Pacinian lamellar cells (p=0.111) and increasing τ_decay_ in Meissner cells (p=0.019, Fig. 2J).

Together, these data reveal that lamellar cells of Pacinian and Meissner corpuscles are intrinsically mechanosensitive. The fast activation kinetics of lamellar MA currents, linear voltage dependence, and lack of ion selectivity are consistent with the ion channel-based mechanotransduction mechanism in somatosensory neurons ^26-30^. Interestingly, the decay rates of MA currents in Pacinian lamellar cells are similar to those observed in slowly inactivating neuronal mechanoreceptors, and Meissner lamellar cell decay rates are reminiscent of fast-and intermediate-inactivating mechanoreceptors ^27,31-34^. The significant differences in the rate and voltage dependence of MA current decay between Pacinian and Meissner lamellar cells from duck bill skin indicate that they each express different mechanically-gated ion channels, or the same channels with alternatively modified function.

Given the similarities between lamellar cells and neuronal mechanoreceptors, we wanted to find out if lamellar cells are excitable. We first asked whether they possess voltage-activated conductances by depolarizing and hyperpolarizing their membranes to different test potentials. Such voltage stimulation of Pacinian lamellar cells failed to reveal voltage-activated potassium, sodium or calcium currents (Extended Data Fig. 2A and B). Moreover, depolarizing current injection failed to evoke any action potentials and instead induced a linear depolarization of the membrane with a slope averaging 2.7 mV/pA, typical of non-excitable cells (Extended Data Fig. 2C). In contrast, lamellar cells from Meissner corpuscles displayed robust voltage-gated potassium currents (Fig. 3A). When these currents were blocked by replacing K^+^ with Cs^+^ in the patch pipette, we identified voltage-gated inward currents that were largely blocked by Cd^2+^ or depletion of extracellular Ca^2+^, suggesting they were mediated by voltage-gated calcium (Ca_v_) channels (Fig. 3B-F). Ratiometric live-cell calcium imaging of duck bill skin revealed that high extracellular potassium-induced depolarization evoked an increase in intracellular calcium in lamellar cells of Meissner, but not Pacinian, corpuscles (Fig. 3G-I), corroborating our finding that Meissner lamellar cells express Ca_v_ channels.

**Fig. 3.**
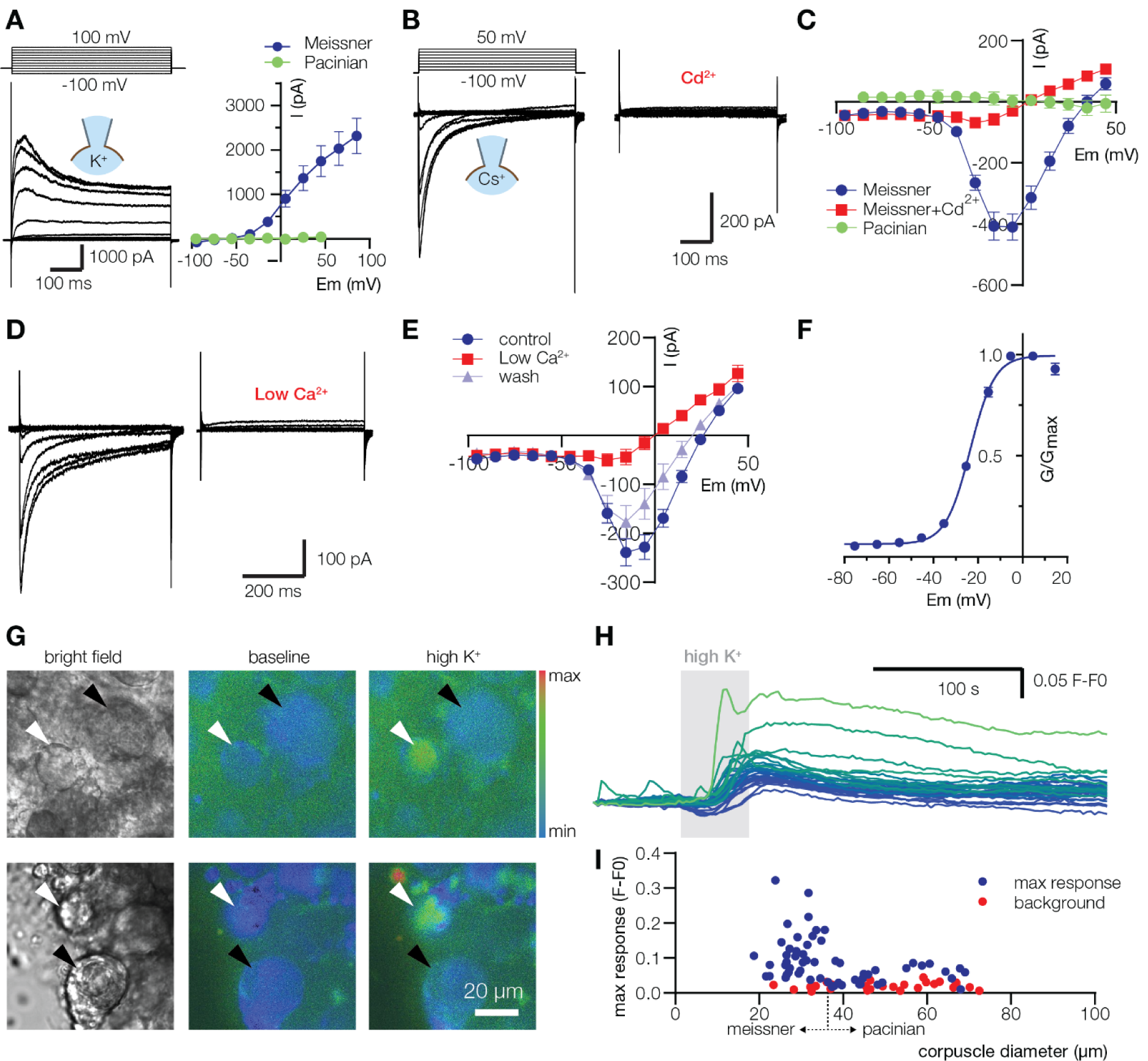
Lamellar cells from Meissner corpuscles express voltage-activated channels. **(A)** Current traces and IV plots of voltage-activated K^+^ currents (mean ± s.e.m., n=12 Meissner and 5 Pacinian lamellar cells). **(B-E)** Current traces and IV plots of voltage-activated Ca^2+^ currents in the presence of pan-Ca_v_ channel blocker 300 µM Cd^2+^ (*B,C*, n=5 cells) and upon depletion of extracellular Ca^2+^ to 20 µM, Low Ca^2+^ (*D, E*, n=7 Meissner and 7 Pacinian lamellar cells). Data are mean ± s.e.m. **(F)** Conductance-voltage relationship of Ca_v_ current, fitted to the Boltzmann equation, with half-maximal activation voltage (V_1/2_) of −23.5 ± 0.4 mV (mean ± s.e.m., n=12 Meissner lamellar cells). **(G)** Representative partial fields of view of live-cell ratiometric Fura-2AM calcium imaging of Meissner (white arrowheads) and Pacinian (black arrowheads) corpuscles in duck bill skin. Application of 135 mM extracellular potassium (high K^+^) elevates intracellular calcium in lamellar cells of Meissner, but not in Pacinian corpuscles or in the neuronal ending within the corpuscles **(H)** Example traces from Meissner corpuscles in response to application of high K^+^. Colors of the traces correspond to the color scale bar in (*G*) based on peak response value. **(I)** Quantification of peak calcium signal in Pacinian and Meissner corpuscles, and in skin areas of comparable sizes devoid of corpuscles (background) in response to high K^+^. Dots represent individual data points. All data are from at least two independent skin preparations.

Having established that Meissner lamellar cells express voltage-gated ion channels, we asked whether they can fire action potentials. Depolarizing current injection triggered repetitive action potential firing in Meissner lamellar cells with a rheobase averaging 16.07 ± 1.9 pA (Fig. 4A and Extended Data Fig. 3A). The voltage-current relationship was strongly rectifying – a characteristic of excitable cells (Extended Data Fig. 3B). In agreement with our finding that Meissner lamellar cells express Ca_v_ channels, the depletion of extracellular Ca^2+^ or addition of Cd^2+^ dampened firing (Fig. 4A, B and E), whereas tetrodotoxin, a blocker of voltage-gated sodium channels, did not (Fig. 4C and E). Transcriptomic analysis revealed that several types of Ca_v_ channel alpha subunits were expressed in duck bill skin (Fig. 4F). However, pharmacological blockade of L-, N, T-, and P/Q-type Ca_v_ channels failed to affect firing (Fig. 4E and S4). In contrast, SNX-482, a specific blocker of R-type (Ca_v_2.3) channels, completely abolished action potential firing (Fig. 4D and E). Thus, action potential firing in Meissner lamellar cells must be mediated by R-type Ca_v_ channels.

**Fig. 4.**
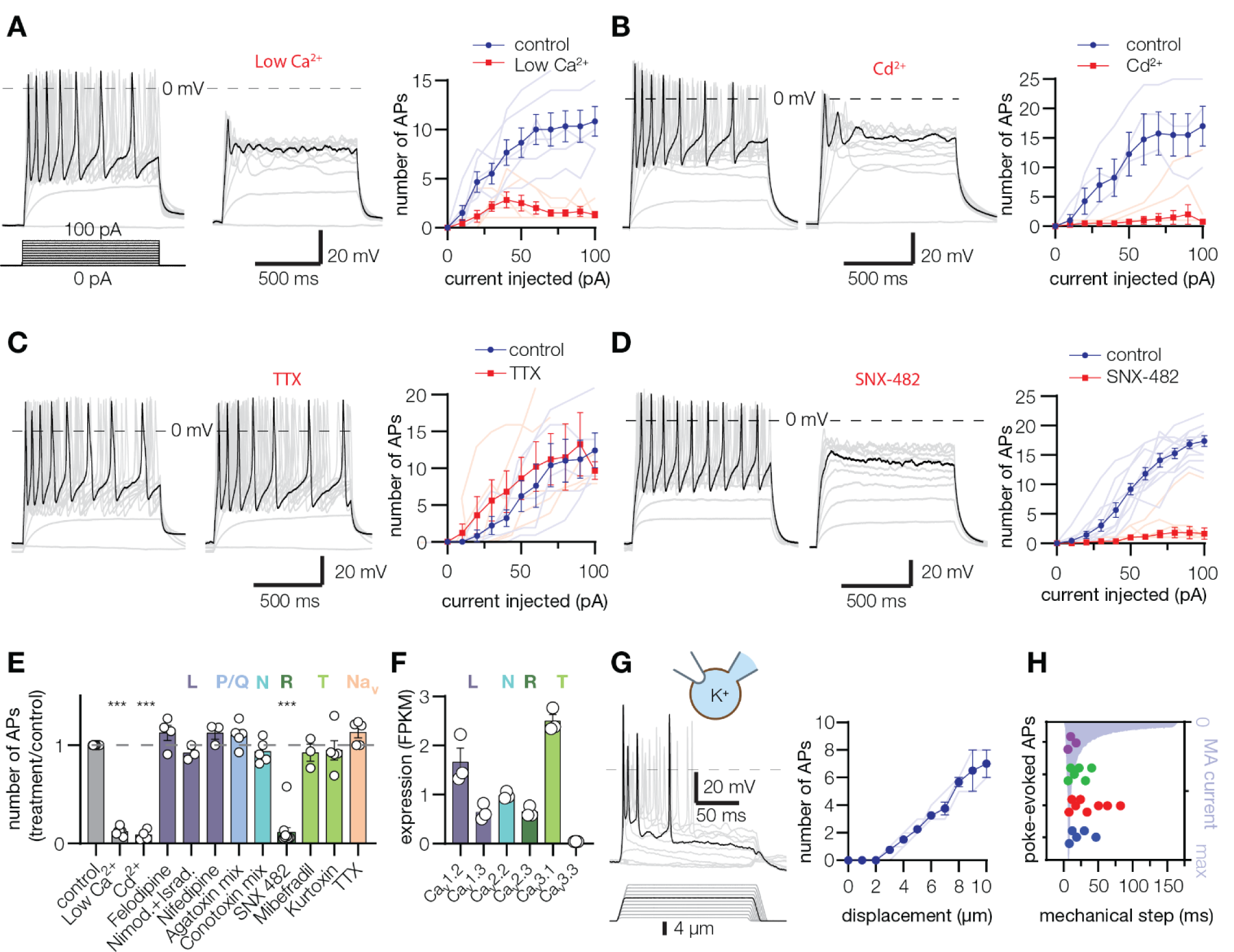
Lamellar cells from Meissner corpuscles are excitable mechanosensors. **(A-D)** Exemplar action potentials (*left, middle* panels) and quantification of spikes (*right* panels) obtained by current injection into Meissner lamellar cells. Firing is inhibited upon depletion of extracellular Ca^2+^ to 20µM, Low Ca^2+^ (*A*, n=6 cells), in the presence of pan-Ca_v_ channel blocker 300 µM Cd^2+^ (*B*, n=4 cells) and R-type Ca_v_2.3 channel blocker 1 µM SNX-482 (*D*, n=11 cells) but not by the voltage-gated sodium channel blocker 100 µM tetrodotoxin, TTX (*C*, n=5 cells). Thin lines in quantification panels represent individual cells, thick lines connect means ± s.e.m. **(E)** Pharmacological profile of Meissner lamellar cell firing in response to a 100 pA current injection, normalized to control treatment. Letters indicate Ca_v_ type selectivity. Na_v_, voltage-gated sodium channel. Data are means ± s.e.m. from at least two independent experiments. Open circles represent individual cells. The effect of treatment is significant, *F*_11,53_=75.57, *p*<0.0001, one-way ANOVA; \*\*\**p*<0.0001 *vs*. control, Dunnett’s post-hoc test. **(F)** Quantification of Ca_v_ channel alpha subunit mRNA expression from duck bill skin, presented as the mean of the number of mRNA fragments per kilobase of exon per million fragments mapped (FPKM) ± s.e.m. Open circles represent samples from individual animals. **(G)** Mechanical stimulation evokes action potential firing in Meissner lamellar cells. Shown are exemplar action potential traces (*left* panel), and quantification of the number of action potentials in response to 150 ms long mechanical stimulation (*right* panel), pooled from three Meissner lamellar cells (thin lines). Thick line connects data means ± s.e.m. **(H)** The number of mechanically-evoked action potentials is maximal when MA current is at its peak. Shown is quantification of the number of action potentials (dots) upon mechanical stimulation of 4 Meissner lamellar cells to 8 µm depth, plotted against peak-normalized MA current profile.

Because the rheobase for Meissner lamellar cell firing was comparable to the amplitude of MA current produced by direct mechanical stimulation, we wondered whether mechanical stimulation alone could elicit firing. Indeed, indentation with a glass probe triggered repetitive firing in Meissner lamellar cells with a threshold of 4.6 ± 0.4 µm (n=7 cells, Extended Data Fig. 3C); the number of action potentials increased in proportion to the degree of indentation (Fig. 4G). Notably, the duration of mechanically-evoked action potentials had the same timing as the duration of the mechanically-evoked current, further supporting the causative relationship between these events (Fig. 4H). Together, these data demonstrate robust mechanically-evoked excitability in Meissner lamellar cells.

We have shown that Meissner lamellar cells are non-neuronal mechanosensors that can generate Ca^2+^-dependent action potentials via R-type Ca_v_ channels. To our knowledge, this is the only non-neuronal cell type that utilizes R-type Ca_v_ channels for firing. We detected mechanosensitivity, but not excitability, in Pacinian outer core lamellar cells. Nevertheless, the exceptionally high input resistance of these cells together with their robust MA currents is sufficient to produce strong touch-induced depolarization without the need for amplification via voltage-gated machinery. The MA currents produced by Pacinian and Meissner lamellar cells are different from each other and from MA currents produced by Piezo2; a mechanically-gated ion channel with a prominent role in somatosensory mechanotransduction in vertebrates ^19,30,35-41^. Whether lamellar MA currents are mediated by Piezo2 with modified function ^42-44^, or by other proteins ^45-47^, remains to be determined.

The identification of active touch detection in lamellar cells within Pacinian and Meissner corpuscles suggests that their function extends beyond passive structural support for the neuronal afferent. That removal of the layers surrounding the afferent ending in Pacinian corpuscles converts neuronal firing from rapidly to slowly adapting has long served as evidence that lamellar cells form a passive mechanical filter that prevent static stimuli from reaching the afferent ^13^. By inference, a similar role has been attributed to the interdigitating protrusions formed between lamellar cells and the neuron in Meissner corpuscles. Although duck Meissner corpuscles display rapidly adapting firing like their mammalian counterparts and have similar frequency tuning characteristics, their lamellar cells form only minimal interdigitations with the neuron. This suggests that extensive mechanical layers around the neuron may be important, but not be the only prerequisite for rapid adaptation. We instead propose that lamellar cells play an active role in shaping the rapid adaptation of afferent firing in response to static stimulation; a process that endows layered and non-layered corpuscles with exquisite sensitivity to transient pressure and vibration. Both types of corpuscle contain molecular components of synaptic machinery ^14-16^, raising the possibility that lamellar cells may shape afferent responses via a synapse-like mechanism.

## METHODS

### Animals

Experiments with Pekin duck embryos (*Anas platyrhynchos domesticus*) were approved by and performed in accordance with guidelines of Institutional Animal Case and Use Committee of Yale University (protocol 2018-11526).

### Preparation of duck bill skin

Pacinian and Meissner corpuscles acquire functionality several days before hatching, and become capable of producing a rapidly adapting discharge in the innervating mechanoreceptor in response to touch as early as E24-26, similar to corpuscles from adult animals ^19-21^. A patch of skin (∼5mm x 10mm) from E24-26 duck embryo was peeled from the dorsal surface of the upper bill, and the epidermis was mechanically removed to expose Pacinian and Meissner corpuscles. Skin was incubated in 2 mg/ml Collagenase P (Roche) in *Krebs* solution (in mM: 117 NaCl, 3.5 KCl, 2.5 CaCl_2_, 1.2 MgCl_2_, 1.2 NaH_2_PO_4_, 25 NaHCO_3_, 11 glucose, saturated with 95% O_2_ and 5% CO_2_ to pH 7.3-7.4 at 22°C) for 20-25 min, washed three times with *Krebs* and imaged external side up on an Olympus BX51-WI upright microscope equipped with an Orca flash 2.8 camera (Hamamatsu).

### Patch-clamp electrophysiology of lamellar cells

Recordings were carried out at room temperature using a MultiClamp 700B amplifier and digitized using a Digidata 1550 (Molecular Devices). Patch pipettes were pulled using a P-1000 puller (Sutter Instruments) from 1.5 mm borosilicate glass with a tip resistance of 1.5-3 MΩ.

Voltage-clamp recordings were acquired in the whole-cell mode using pClamp 10 software, sampled at 20 kHz and low-pass filtered at 10 kHz. Voltage-clamp experiments were recorded from a holding potential of −80 mV, using the following solutions (in mM). *Internal-Cs:* 133 CsCl, 5 EGTA, 1 CaCl_2_, 1 MgCl_2_, 10 HEPES, 4 Mg-ATP, 0.4 Na_2_-GTP pH 7.3 with CsOH. *Internal-K:* 135 K-gluconate, 5 KCl, 0.5 CaCl_2_, 2 MgCl_2_, 5 EGTA, 5 HEPES, 5 Na_2_ATP and 0.5 GTP-TRIS pH 7.3 with KOH. *Bath Ringer:* 140 NaCl, 5 KCl, 10 HEPES, 2.5 CaCl_2_, 1 MgCl2, 10 glucose, pH 7.4 with NaOH. Voltage-gated potassium currents were recorded using *Internal-K* and *Bath Ringer*. Currents were elicited by 500 ms voltage steps from −100 mV, in 10 mV increments. Voltage-gated sodium and calcium (Ca_v_) currents were recorded using *Internal-Cs* and *Bath Ringer* supplemented or not with 300 µM CdCl_2_ or 20 µM CaCl_2_. Currents were elicited using 500 ms voltage steps from −100 mV, in 10 mV increments. Each voltage step was proceeded by a 500 ms hyperpolarizing step to −120 mV to remove channel inactivation. Leak current was subtracted using the P/4 protocol. Series resistance was compensated at 50%. Peak Ca_v_ currents were converted to conductance using the equation *G = I / (V*_*m*_ *– E*_*rev*_), where *G* is the conductance, *I* is the peak Ca_v_ current, *V*_*m*_ is the membrane potential and *E*_*rev*_ is the reversal potential. The conductance data were fit with the modified Boltzmann equation, *G = G*_*min*_ *+ (G*_*max*_ *– G*_*min*_*) / (1 + exp^([V*_*1/2*_ *– V*_*m*_*]/k))*, where *G*_*min*_ and *G*_*max*_ are minimal and maximal conductance, respectively, *V*_*m*_ is the voltage, *V*_*1/2*_ is the voltage at which the channels reached 50% of their maximal conductance, and *k* is the slope of the curve.

Mechanically-activated currents were recorded in *Internal-Cs* and *Bath Ringer* at a −60mV holding potential. After whole cell formation, a blunt glass probe (2-4 µm at the tip) mounted on a piezoelectric driven actuator (Physik Instrumente GmbH) was positioned to touch the corpuscle at the side opposite to the patch pipette. The probe mounted was moved at a velocity of 800 μm/s toward the corpuscle in 1-μm increments, held in position for 150 ms and then retracted at the same velocity.

To visualize lamellar cells, Lucifer Yellow was added to internal solution at concentration of 2 mg/ml. Resting membrane potentials were measured upon break-in using *Internal-K* and *Bath-Ringer*. Voltage-clamp experiments and resting membrane potential measurements were corrected offline for liquid junction potential calculated in Clampex 10.7.

Current-clamp experiments were recorded using *Internal-K* and *Krebs* in the bath. Recordings were started 2 minutes after break-in to stabilize the action potential firing. Changes in membrane potential were recorded in response to 1 s current pulses from a 0 to −30 pA holding, in 10 pA increments. Current-clamp experiments were not corrected for liquid junction potential. For pharmacological experiments, bath solution was supplemented with the following: 300 µM CdCl_2_, 20 µM CaCl_2_, 10 µM Felodipine (Abcam), a mix of 10 µM Nimodipine and 5 µM Isradipine (Alomone), 10 µM Nifedipine (Alomone), Agatoxin mix (1 µM ω-Agatoxin IVA and 1 µM ω-Agatoxin TK from Alomone), Conotoxin mix (5 µM ω-Conotoxin CnVIIA, 10 nM ω-Conotoxin CVIB, 10 nM ω-Conotoxin CVIE, 1 µM ω-Conotoxin MVIIC and 1 µM ω-Conotoxin MVIID, from Alomone), 1 µM SNX-482 (from Alomone or Peptides International), 5 µM Mibefradil*2HCl, 200 nM Kurtoxin (Alomone), 200 µM Tetrodotoxin citrate (Tocris). Paired recordings were performed 1-10 min after the addition of small molecule drugs, or 1-20 min after the addition of peptide toxins.

### Preparation of trigeminal neurons

Trigeminal neurons from embryonic duck (E24-E26) were acutely dissociated as previously described ^19,25^. Dissected duck TG were chopped with scissors in 500 μl ice-cold HBSS, dissociated by adding 500 μl of 2 mg/ml collagenase P (Roche) dissolved in HBSS and incubated for 15 min at 37 °C, followed by incubation in 500 μl 0.25% trypsin-EDTA for 10 min at 37 °C. The trypsin was then removed and the residual trypsin was quenched by adding 750 μl pre-warmed DMEM+ medium (DMEM supplemented with 10% FBS, 1% penicillin/streptomycin and 2 mM glutamine). Cells were triturated gently with plastic P1000 and P200 pipettes and collected by centrifugation for 3 min at 100 × g. Cells were resuspended in DMEM+ medium and plated onto the Matrigel (BD Bioscience, Billerica, MA) -precoated coverslips in a 12-well cell-culture plate. 0.5 ml DMEM+ medium was added into each well following incubation at 37 °C in 5% CO_2_ for 30-45 min. MA current measurements were performed within 48 hours after plaiting.

### Patch-clamp electrophysiology of trigeminal neurons

Voltage-clamp recordings were acquired in the whole-cell mode using pClamp software using 1.5 mm borosilicate glass with a tip resistance of 1.5-5 MΩ. Recordings were performed in *Bath Ringer*, sampled at 20 kHz and low-pass filtered at 2-10 kHz. Internal solution contained (in mM): 130 K-methanesulfonate, 20 KCl, 1 MgCl_2_, 10 HEPES, 3 Na_2_ATP, 0.06 Na_2_GTP, 0.2 EGTA, pH 7.3, with KOH (final [K^+^] = 150.5 mM). Prior to mechanical stimulation, current was injected in current-clamp mode to elicit neuronal firing. Mechanical stimulation was performed using a blunt glass probe positioned at 32° −55° relative to the cell as described above for corpuscles. Membrane potential was clamped at −60 mV. Neurons with MA current were classified based on the rate of MA current inactivation (τ_inact_) as fast inactivating (τ_inact_ < 10 ms), intermediately inactivating (τ_inact_ = 10-30 ms) and slow inactivating (τ_inact_ > 30 ms) as previously described ^25^: the decaying component of MA current was fit to the single-exponential decay equation: *I=ΔI*exp^(-t/τ*_*inact*_), where *ΔI* is the difference between peak MA current and baseline, *t* is the time from the peak current (the start of the fit), and τ_inact_ is the inactivation rate. Resultant τ_inact_ for each neuron represent an average from traces with the top 75% of MA amplitude ^30^. Mechanically activated current rise (τ_rise_) time was quantified by fitting a single-exponential function in similar manner as for τ_inact_.

### RNA Sequencing

Total RNA was isolated from duck bill skin using the TRIzol reagent (ThermoFisher, Waltham, MA) according to manufacturer’s instructions. RNA integrity was assessed based on RIN values obtained with Agilent Bioanalyzer. Library preparation and sequencing were carried out at the Yale Center for Genome Analysis. mRNA was purified from ∼200 ng total RNA with oligo-dT beads. Strand-specific sequencing libraries were prepared using the KAPA mRNA Hyper Prep kit (Roche Sequencing, Pleasanton, CA). Libraries were sequenced on Illumina NovaSeq sequencer in the 100 bp paired-end sequencing mode according to manufacturer’s protocols with multiple samples pooled per lane. A total of ∼50-69 million sequencing read pairs per sample were obtained. The sequencing data was processed on the Yale High Performance Computing cluster. Raw sequencing reads were filtered and trimmed to retain high-quality reads using Trimmomatic v0.36 with default parameters. Filtered high-quality reads from all samples were aligned to duck reference genome using the STAR aligner v2.5.4b with default parameters. The reference genome (Anas platyrhynchos, BGI_duck_1.0) and gene annotation (NCBI Release 102) were obtained from the National Center for Biotechnology Information (accessed on 8/5/2018). The gene annotation was filtered to include only protein-coding genes. Aligned reads were counted by featureCounts program within the Subread package v1.6.2 with default parameters. Raw read counts were processed and converted to ‘‘mRNA fragments per kilobase of exon per million mapped fragments’’ (FPKM) values by EdgeR v3.22.3. The RNA sequencing data was deposited to the Gene Expression Omnibus, accession number: GSE155529.

### Calcium Imaging

Live-cell ratiometric calcium imaging was performed on duck bill skin patches at room temperature using Axio-Observer Z1 inverted microscope (Zeiss) equipped with an Orca-Flash 4.0 camera (Hamamatsu) using MetaFluor software (Molecular Devices). After collagenase treatment, skin patch was loaded with 10 mM Fura 2-AM (Thermo Fisher) and 0.02% Pluronic F-127 in Ringer solution for 30 min at room temperature and washed 3 times with *Ringer* solution. The skin was then visualized and exposed to a *high-K*^*+*^ solution, containing (in mM): 10 NaCl, 135 KCl, 2 CaCl_2_, 2 MgCl_2_ and 10 glucose, 10 HEPES pH 7.4 (with KOH). Background signal was quantified from skin areas devoid of corpuscles.

### Electron microscopy

Freshly peeled duck bill skin was fixed in Karnovsky fixative at 4°C for one hour, washed in 0.1M sodium cacodylate buffer pH 7.4, post-fixed in 1% osmium tetroxide for one hour in the dark on ice. The tissue was stained in Kellenberger solution for one hour at room temperature after washing in distilled water, dehydrated in a series of alcohols and propylene oxide then embedded in Embed 812 and polymerized overnight at 60°C. All solutions were supplied by Electron Microscopy Sciences Hatfield, PA. Ultrathin sections were obtained on a Leica Ultracut UCT ultramicrotome at 70 nm, stained in 1.5% aqueous uranyl acetate and Reynolds Lead stains and imaged on a FEI Tecnai G2 Spirit BioTWIN electron microscope.

### Quantification and statistical analysis

Electrophysiological data from corpuscles and trigeminal neurons were obtained from skin preparations from at least three animals. All measurements were taken from distinct samples. Data were analyzed and plotted using GraphPad Prism 8.4.3 (GraphPad Software Inc) and expressed as means ± s.e.m. or as individual points. Statistical tests were chosen based on experimental setup, sample size and normality of distribution, as determined by the Kolmogorov-Smirnov test, and are specified in figure legends. Adjustments for multiple comparisons were performed where appropriate.

### Data availability

The RNA sequencing data was deposited to the Gene Expression Omnibus, accession number GSE155529. Other data are available from the corresponding authors upon request.

## Acknowledgements

We thank Ever Schneider for performing pilot studies, SueAnn Mentone for electron microscopy imaging, Ruslan Dashkin for help with data visualization, and members of the Bagriantsev and Gracheva laboratories for their contributions throughout the project.

## Author contributions

E.O.G. and S.N.B. conceived the project. Y.A.N. and E.O.G. developed the skin preparation. Y.A.N. performed electrophysiological and calcium imaging recordings from corpuscles. V.V.F. performed transcriptomic analysis. E.O.A. performed electrophysiological recordings from trigeminal neurons. Y.A.N., E.O.G. and S.N.B. wrote the paper.

---

**Extended Data Fig. 1.**
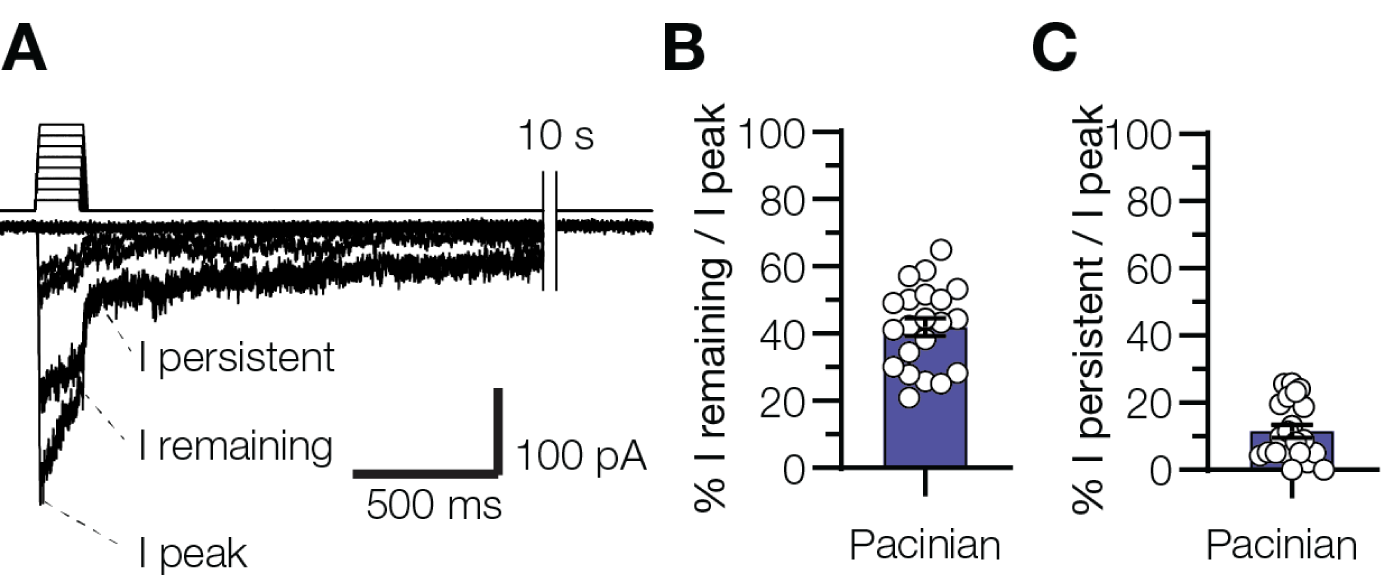
Mechanically-activated currents in lamellar cells within Pacinian corpuscles. **(A)** Exemplar MA current traces from a Pacinian lamellar cell showing the decay of MA current to baseline. **(B, C)** Quantification of MA current amplitude in Pacinian lamellar cells immediately before (*B*) and 10 ms after retraction of the probe (*C*) relative to peak MA current amplitude. Data are means ± s.e.m. from at least three independent skin preparations. Open circles denote individual cells.

**Extended Data Fig. 2.**
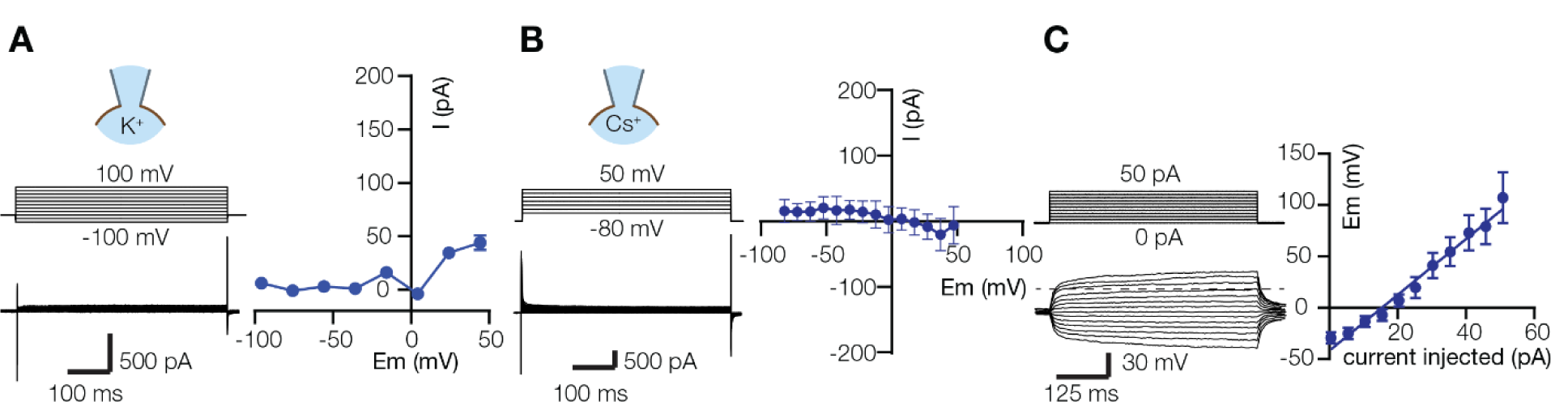
Lamellar cells from Pacinian corpuscles lack voltage-gated currents. **(A, B)** Exemplar current-voltage relationships recorded in response to voltage steps with K^+^-based (*A*) or Cs^+^-based (*B*) internal solution. Data are mean ± s.e.m. from 5 and 7 Pacinian lamellar cells, respectively. In *A*, the error bars are smaller than the symbols. **(C)** Exemplar voltage traces in Pacinian lamellar cells and quantification of membrane potential change in response to current injection, fitted to the linear equation (n=7 cells). Data are means ± s.e.m., collected from at least two independent skin preparations.

**Extended Data Fig. 3.**
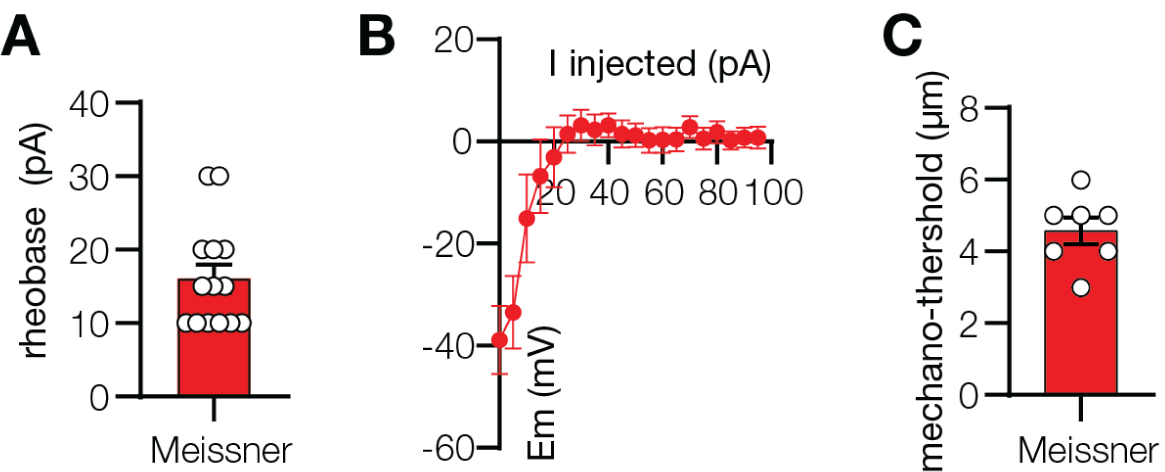
Lamellar cells from Meissner corpuscles are excitable. **(A)** Quantification of Meissner lamellar cell firing threshold in response to current injection. Data are means ± s.e.m. Each dot represents an individual cell. **(B)** Quantification of peak membrane potential of Meissner lamellar cells in response to current injection. Data are presented as means ± s.e.m. from 8 individual cells. **(C)** Quantification of action potential firing threshold evoked in Meissner lamellar cells by mechanical indentation. Data are means ± s.e.m. Each dot represents an individual cell.

**Extended Data Fig. 4.**
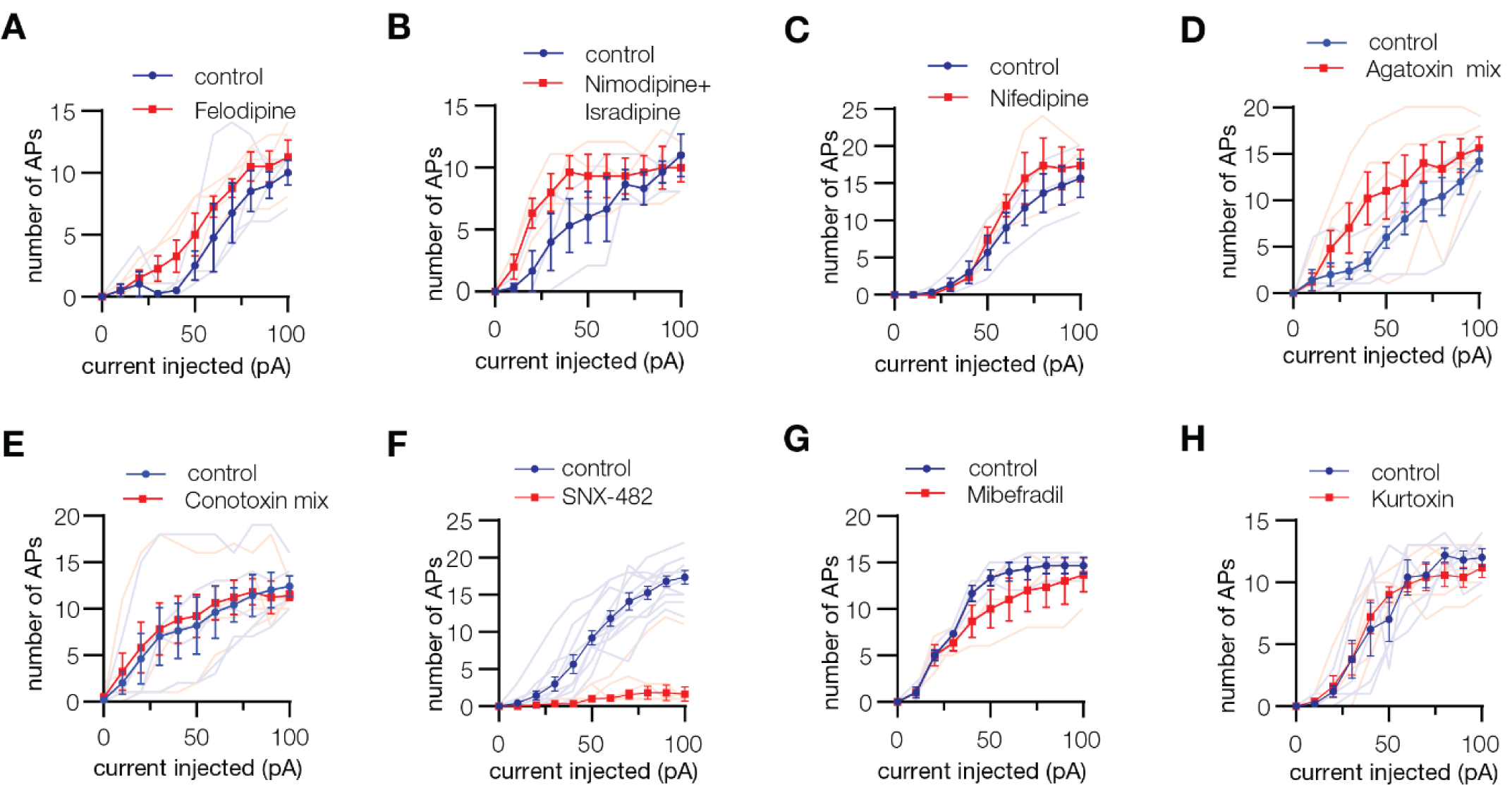
Pharmacological profile of Meissner lamellar cell firing. Quantification of the number of action potentials in response to current injection in the presence of indicated pharmacological agents: 10 µM Felodipine, a mix of 10 µM Nimodipine and 5 µM Isradipine, 10 µM Nifedipine, Agatoxin mix (1 µM ω-Agatoxin IVA and 1 µM ω-Agatoxin TK), Conotoxin mix (5 µM ω-Conotoxin CnVIIA, 10 nM ω-Conotoxin CVIB, 10 nM ω-Conotoxin CVIE, 1 µM ω-Conotoxin MVIIC and 1 µM ω-Conotoxin MVIID), 1 µM SNX-482, 5 µM Mibefradil, 200 nM Kurtoxin. Thin lines represent individual cells, thick lines connect means ± s.e.m. Data were obtained from at least two independent experiments.

